# Biological variation in the sizes, shapes and locations of visual cortical areas in the mouse

**DOI:** 10.1101/414698

**Authors:** Jack Waters, Eric Lee, Nathalie Gaudreault, Fiona Griffin, Jerome Lecoq, Cliff Slaughterbeck, David Sullivan, Colin Farrell, Jed Perkins, David Reid, David Feng, Nile Graddis, Marina Garrett, Yang Li, Fuhui Long, Chris Mochizuki, Kate Roll, Jun Zhuang, Carol Thompson

## Abstract

Visual cortex is organized into discrete sub-regions or areas that are arranged into a hierarchy and serve different functions in the processing of visual information. In our previous work, we noted that retinotopic maps of cortical visual areas differed between mice, but did not quantify these differences or determine the relative contributions of biological variation and measurement noise. Here we quantify the biological variation in the size, shape and locations of 11 visual areas in the mouse. We find that there is substantial biological variation in the sizes of visual areas, with some visual areas varying in size by two-fold across the population of mice.

## INTRODUCTION

Mammalian neocortex is generally considered to be organized into discrete anatomically and functionally defined sub-regions or areas. For example, a major portion of posterior cortex is concerned primarily with vision and is divided into discrete visual areas, >20 areas in primates (Gattass *et al.*, 2005) and ∼15 in the mouse (Zhuang et al., 2017). Visual areas are arranged into a hierarchy and serve different functions in the processing of visual information (Van Essen & Maunsell, 1983) and yet the sizes and shapes of visual areas appear to vary across individuals of a given species. In mice for example, maps of visual cortex consist of a stereotyped collection of visual areas, in approximately the same relative locations in each mouse, but the relative positions of areas and in their sizes and shapes appear to differ substantially between mice. Such differences in the organization of cortex might affect the processing of visual information across mice. How much variation is there in the locations, sizes and shapes of cortical areas from mouse to mouse?

Only a few studies describe differences in the size, shape and locations of cortical areas across a population of individuals. Differences have been described in the size and shape of primary visual cortex in humans, macaques, cats, rats and mice, the consensus being that V1 differs in size more than in shape (Van Essen, Newsome & Maunsell, 1984; Duffy, Murphy & Jones, 1998; Hinds *et al.*, 2008; Garrett *et al*., 2014). Necessarily, comparisons were across small numbers (<25) of individuals and no attempt was made to determine whether the differences resulted from differing organization of cortex across individuals (biological variation) or simply measurement error (noise). Separating biological variation from measurement noise requires statistical analysis of measurements from large numbers of animals. To our knowledge, nobody has assessed biological variation (after separation from measurement noise) of any cortical area in any species.

In our previous work, we noted that retinotopic maps of cortical visual areas differed between mice (Zhuang *et al.*, 2017), but did not quantify these differences or determine the relative contributions of biological variation and measurement noise. Here we quantify the variability in size, shape and locations of cortical visual areas in the mouse

## RESULTS AND DISCUSSION

Retinotopic maps differ across mice (Figure 1 - figure supplement 1). The differences between maps presumably include biological variation and inaccuracies, or measurement noise in the mapping process. Under the assumption that the retinotopic map is invariant in an individual, repeated map generation from a mouse will result in maps that differ only because of measurement noise. This assumption provides a method to isolate biological variation, given maps from a large enough collection of mice and two or more maps from each mouse. From such a data set, one can estimate (1) the effects of measurement noise, by comparing retinotopic maps across mapping sessions in each mouse, and (2) the combined effects of biological variability and measurement noise, by comparing retinotopic maps across mice. A second assumption, that measurement noise and biological variation are independent, permits the isolation of biological variation by subtraction of the variance of the between-session comparisons from the variance of the between-mouse comparisons. Implicitly, a third assumption is made: that there are no additional sources of measurement noise when comparing maps across mice, relative to the comparison across mapping sessions.

**Figure 1.**
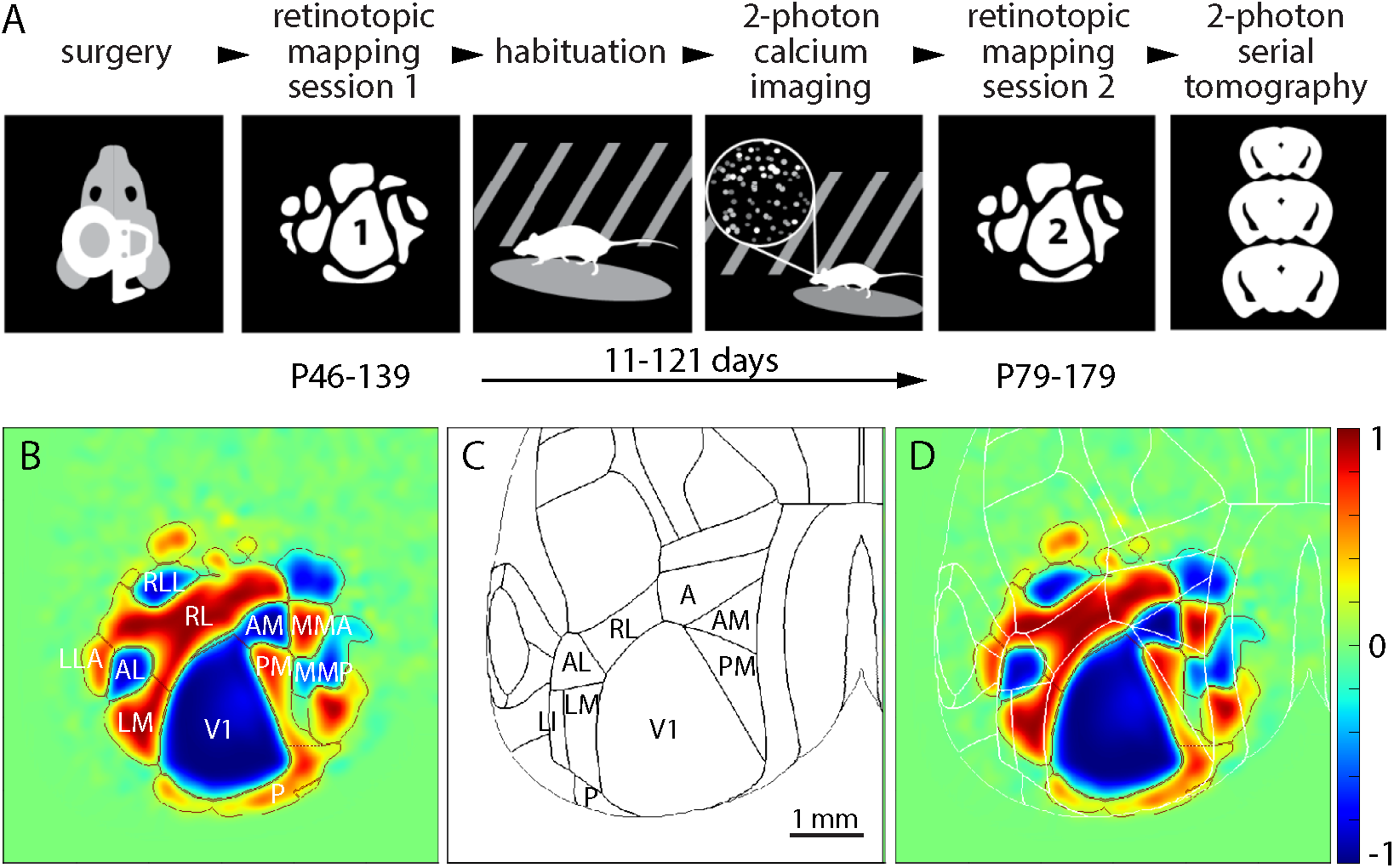
Repeated retinotopic mapping of mouse cortical areas. (A) Workflow of the Allen Brain Observatory, including implantation of a 5 mm diameter cranial window and retinotopic mapping at two time points, separated by 11-121 days. (B) Mean field sign map from 60 mice, with borders and labels for the 11 visual areas studied here. (C) Borders of cortical areas in the 3D Allen Mouse Brain Reference Atlas. Visual areas are labeled. (D) Mean sign map aligned to the 3D Allen Mouse Brain Reference Atlas, with atlas borders in white. The color scale bar applies to all field sign maps throughout the paper.

We generated retinotopic maps for 60 adult mice by intrinsic signal imaging (Schuett *et al.*, 2002; Kalatsky & Stryker, 2003; Marshel *et al.*, 2011). Each mouse was imaged twice, 11-121 days apart (figure 1A). As in our previous publication (Zhuang *et al.*, 2017), retinotopy was displayed using the visual field sign (Garrett *et al.*, 2014) and field sign maps were segmented into retinotopically-defined cortical areas using a numerical routine (Juavinett *et al.*, 2017; Zhuang *et al.*, 2018; Materials and Methods). Maps were aligned to the 3D Allen Mouse Brain Reference Atlas, assisted by surface vasculature images acquired during mapping (Figure 1 - figure supplement 2).

The mean sign map included 11 visual areas (figure 1B; V1, RL, LM, AM, PM, P, RLL, AL, LLA, MMA, MMP). Consistent with our previous paper (Zhuang *et al.*, 2017), there was an almost continuous ring of field sign positive regions around V1 and the lateral retinotopic border of V1 was ∼300 μm medial to the architectonic border (figure 1C, D).

### Biological variation in the locations of visual area

To test for biological variation in the locations of visual areas, we compared maps across mice and across imaging sessions. We reduced each map to a set of points, each point representing the centroid of a field sign patch and made pairwise comparisons of maps. As a measure of the difference between each pair of maps, we calculated the paired patch distance (ppd; figure 2 – figure supplement 1). From 60 mice, two imaging sessions per mouse, we made 60 pairwise comparisons across sessions (one per mouse) and 59 * 60 = 3540 pairwise comparisons across mice. For each pair of maps, we defined paired patch distance as the mean of the distances between corresponding field sign patches:

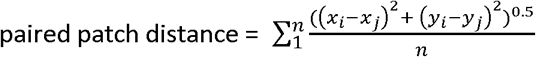

where, x and y are the coordinates of the centroid of each field sign patch in m-l and a-p axes, subscripts i and j denote the two maps being compared, n is the number of field sign patches in both maps

**Figure 2.**
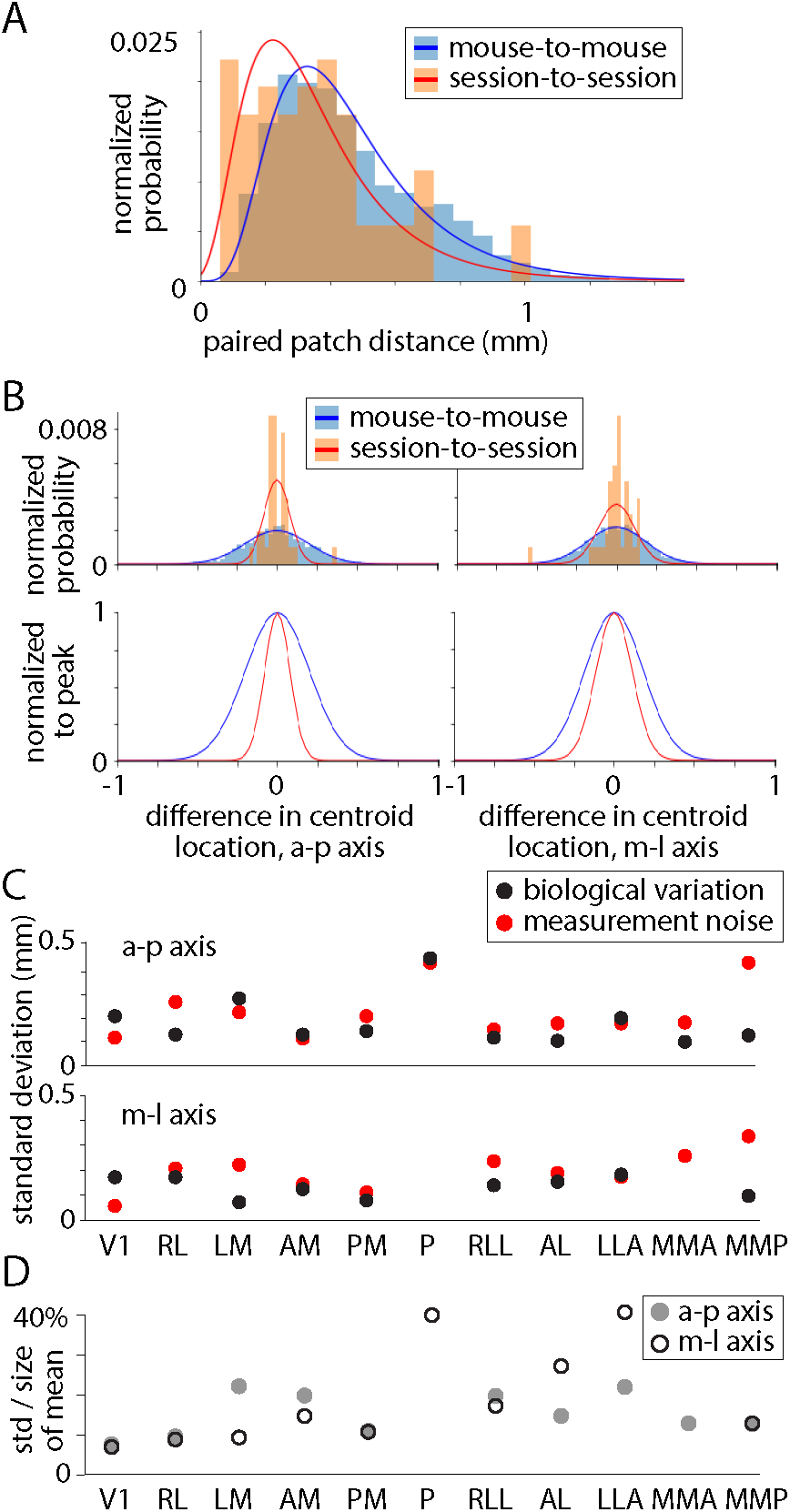
Biological variation in the locations of field sign patches. (A) Frequency histogram of inter-patch distances for 3540 pairwise mouse-to-mouse comparisons and 60 pairwise session-to-session comparisons, after elimination of translation, scale and rotation differences between maps. Frequency is displayed as a fractional probability, the integral of each distribution being 1. Lines are lognormal fits. (B) Frequency histograms illustrating mouse-to-mouse and session-to-session differences in centroid location for AM. Frequency is displayed as a normalized probability, the integral of each distribution being 1. Lines are fits to a normal distribution. Below: same distributions normalized to their peaks. (C) Standard deviations of biological variation and measurement noise for distributions of differences in centroid location. (D) Standard deviation of the biological variation of centroid positions, in a-p and m-l axes, expressed as a percentage of the maximum width of the mean of each patch.

There was no correlation between the time between imaging sessions and session-to-session ppd, consistent with our assertion that maps did not change between imaging sessions (figure 1 – figure supplement 1B). Mouse-to-mouse and session-to-session distributions were significantly different (p = 4.6 x 10^-9^, Mann-Whitney U test), with mouse-to-mouse differences being greater than session-to-session differences. We conclude that there is significant biological variation in the retinotopic map.

For each pairwise comparison of maps, the one map could be translated relative to the other, or rotated, scaled or the relative locations of visual areas might differ. Any of these four parameters might be the source of the observed biological variability. In addition, inaccuracies in our alignment process might give rise to translation and rotation errors that might differ between mouse-to-mouse and session-to-session comparisons and therefore be interpreted, erroneously, as biological variation. To separate differences in translation, rotation, scale and shape (shape, in this context, is the relative locations of patches), we adopted Procrustes superimposition, an analytical approach used to compare shapes in biological populations (Goodall, 1991; Webster & Sheets, 2010; Zelditch, Swiderski & Sheets, 2012). The approach involves the sequential estimation and removal of translation, rotation and scaling to leave differences in only the locations of patches.

We found no compelling evidence for biological variation in the location of V1, or in the size or rotation of retinotopic maps. Across maps, the centroid of V1 moved by up to 1 mm in m-l and a-p axes, but the distributions of mouse-to-mouse and session-to-session differences were similar (p > 0.01, Mann-Whitney U test). As a measure of the scale of each retinotopic map we calculated a statistic we call the ‘centroid size’ (Zelditch, Swiderski & Sheets, 2012) from the centroids of areas V1, RL and PM, all three of which were in every field sign map. Mouse-to-mouse and session-to-session distributions of differences in centroid size were similar (p > 0.01, Mann-Whitney U test). Finally, we observed no difference in rotation of maps (maximum difference in rotation <±0.5 degree; mouse-to-mouse and session-to-session distributions were not significantly different, p = 0.28, Mann-Whitney U test). We aligned maps, eliminating differences in location, scale and rotation, and re-tested for biological variation, finding that a significant difference between the distributions remained (p = 4.4 x 10^-5^, Mann-Whitney U test; figure 2A). We conclude that there is biological variation in the overall shape of the retinotopic map in the mouse, in other words, that the field sign patches are in different locations in different mice.

We further quantified the differences in centroid location for each field sign patch, aiming to calculate the biological variation in location for each patch. Our approach is similar to that of Garrett *et al.* (2014), but here we separate biological variation and measurement noise. Mouse-to-mouse differences in centroid location were greater than session-to-session differences for all 11 patches and in both a-p and m-l axes (figure 2B, C), permitting us to estimate biological variation by subtraction of variances. For most patches, biological variation and measurement noise were approximately equal contributors to mouse-to-mouse differences in the visual area map (figure 2C). The standard deviation of biological variation was ∼200 μm for many patches or ∼15 % of the diameter of the mean patch (figure 2D).

### Size and shape of each field sign patch

When comparing retinotopic maps from different mice, there appears to be striking variation in the sizes and shapes of most patches in the map (figure 1 – figure supplement 1). We further explored biological variation in the size and shape of each field sign patch, to determine whether there is biological variation and to quantify it.

We began by comparing mouse-to-mouse and session-to-session differences in the area of each field sign patch (figure 3A). For 9 of 11 patches, the variance of the mouse-to-mouse distribution was greater than the variance of the session-to-session differences, but the distributions were significantly different only for V1 and not for the remaining 10 patches (p < 0.0045, Levene’s test). For most patches, measurement noise was the main source of variability in patch size, but biological variation was nonetheless substantial at 16-56% of the area of the mean patch (figure 3B). The patch with the least biological variation in size was V1, with a standard deviation of 16%. Hence in 50% of mice, V1 is >11% smaller or larger than average and in 5% of mice, V1 is >32% smaller or larger than average. The standard deviation of biological variation for AM was 33%, approximately equal to the mean biological variation for all patches (32%). In 50% of mice, AM is >22% smaller or larger than average and in 5% of mice, AM is >64% smaller or larger than average. There was no correlation between the area of V1 and the age (figure 3D) or weight (figure 3E) of the mouse and the area of V1 differed little with sex or Cre line (figure 3G). There was no correlation between the sizes of patches across mice (p < 0.05/11, Spearman rank-order correlation), so the sizes of neighboring patches are unrelated.

**Figure 3.**
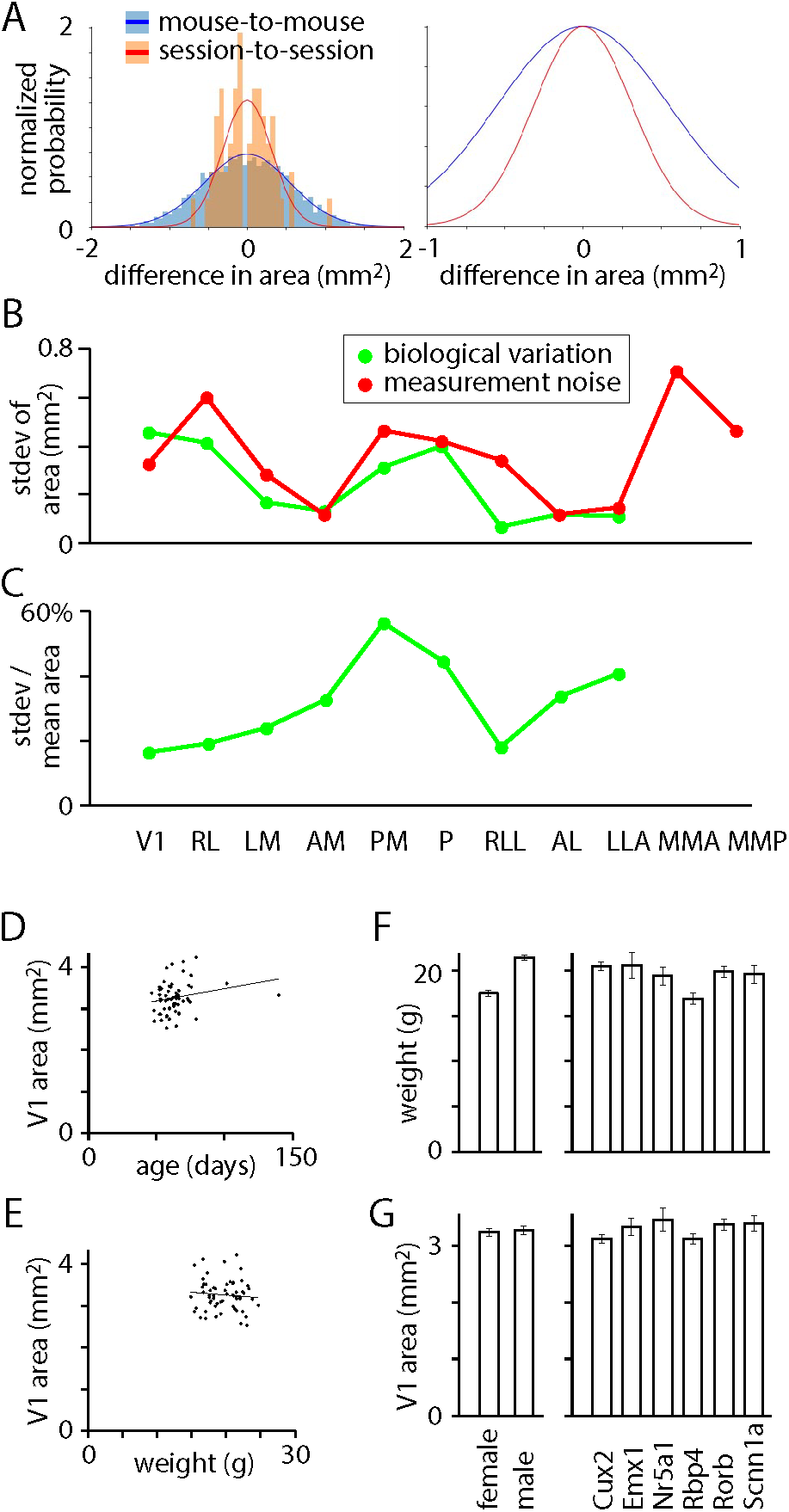
Biological variation in the sizes of field sign patches. (A) Histogram of mouse-to-mouse and session-to-session differences in area of V1. Frequency is displayed as a fractional probability, the integral of each distribution being 1. Lines are normal distributions. Right: same distributions normalized to their peaks. (B) Standard deviations of biological variation and measurement noise for distributions of differences in area. (C) For each patch, standard deviation of biological variation in patch area as a percentage of the area of the mean patch. (D) Area of V1 in the first imaging session, as a function of postnatal age during the first imaging session. Line is a linear fit. (E) Area of V1 in the first imaging session, as a function of weight during the first imaging session. Line is a linear fit. (F) Weights of mice at first imaging session, sorted by sex (left) and Cre line (right). Mean ± SEM. (G) V1 area in first imaging session, sorted by sex (left) and Cre line (right). Mean ± SEM.

We investigated biological variation in the shapes of field sign patches using the Jaccard index, defined as the area of intersection of two patches divided by the area of their union. Jaccard index ranges from 0 (no overlap) to 1 (identical shapes). We calculated differences pairwise for mouse-to-mouse and for session-to-session comparisons, resulting in mouse-to-mouse and session-to-session distributions of the Jaccard index for each patch. The median Jaccard indices of mouse-to-mouse and session-to-session distributions were similar for all patches (figure 3A) and there was no patch with a mouse-to-mouse median greater than its session-to-session median. Consistent with this conclusion, plots of the cumulative mouse-to-mouse and session-to-session differences were similar (figure 3B), with no indication that mouse-to-mouse differences were greater than session-to-session differences in patch shape. Hence we find no evidence for biological variation in the shapes of visual areas, the apparent variability across mice resulting from measurement noise.

We have assumed that biological variation is the only difference between mouse-to-mouse and session-to-session comparisons. Might some other mouse-to-mouse difference have been classified as biological variation? Alignment errors are a concern, but the resulting errors would likely affect the locations of visual areas and have little effect on their sizes. Hence alignment errors are unlikely to account for the observed biological variation in the sizes of visual areas. Mouse-to-mouse differences in the segmentation of field sign patches, arising from differences in the signal-to-noise ratio of the underlying images, might contribute to the biological variation in size of patches. We expect the effect to be greater for peripheral patches (patch P, for example) than for V1 and other patches near the core of the map.

In summary, we assessed biological variation in the sizes, shapes and locations of field sign patches in visual cortex of the mouse. Our results provide no evidence for biological variation in the shapes of visual areas and evidence of only modest biological variation in location, but the sizes of visual areas differed substantially between mice, with the area of the average region varying ∼2-fold across the population of mice. We were surprised by this result. We reasoned that there is some minimum amount of cortical tissue required for each visual area to perform its functions and likely sufficient, but not excess cortex is allocated to each visual area. If the functions of each visual area are invariant across mice, it seems likely that the required volume of cortex for each visual area is similar across mice. Hence our expectation was that there might be variability in the locations and shapes of visual areas, but that size would be more constrained. Our results are not consistent with this expectation, raising the possibility the roles of visual areas in the processing of visual information is more flexible than we had appreciated.

## FIGURE LEGENDS

**Figure 1 - figure supplement 1.**
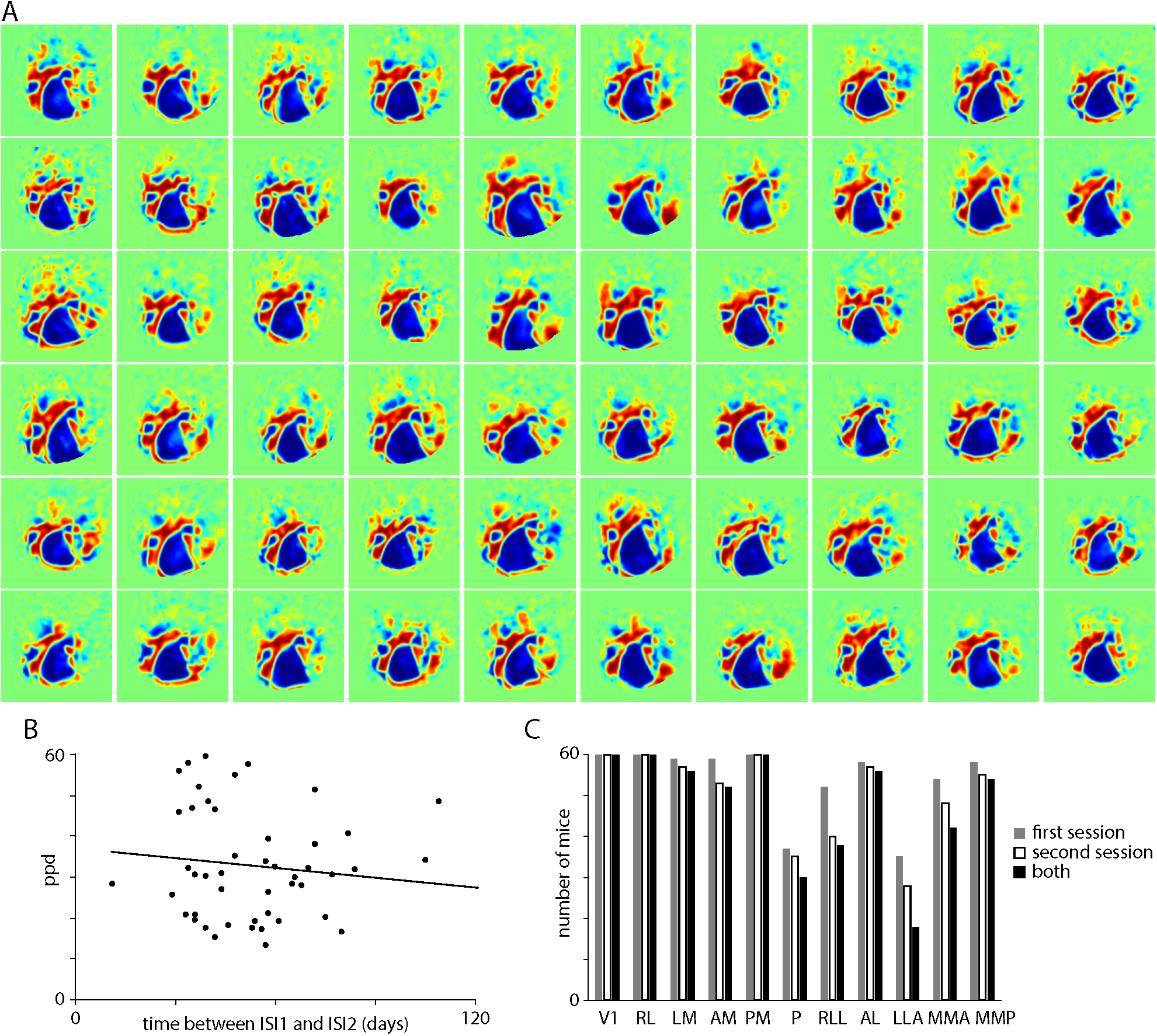
Field sign maps from 60 mice. (A) Field sign maps from the first imaging session for each of 60 mice, to illustrate the mouse-to-mouse variability in field sign maps. (B) Plot of paired patch distance (ppd) as a function of time between ISI1 and ISI2. Line: best linear fit, slope -0.08. (C) Histogram of the number of times each patch appeared in ISI maps. Each patch could occur in a maximum of 60 maps (one for each of 60 mice) for each of the first and second imaging sessions. The right column indicates the number of mice in which the patch was visible in both imaging sessions.

**Figure 1 - figure supplement 2.**
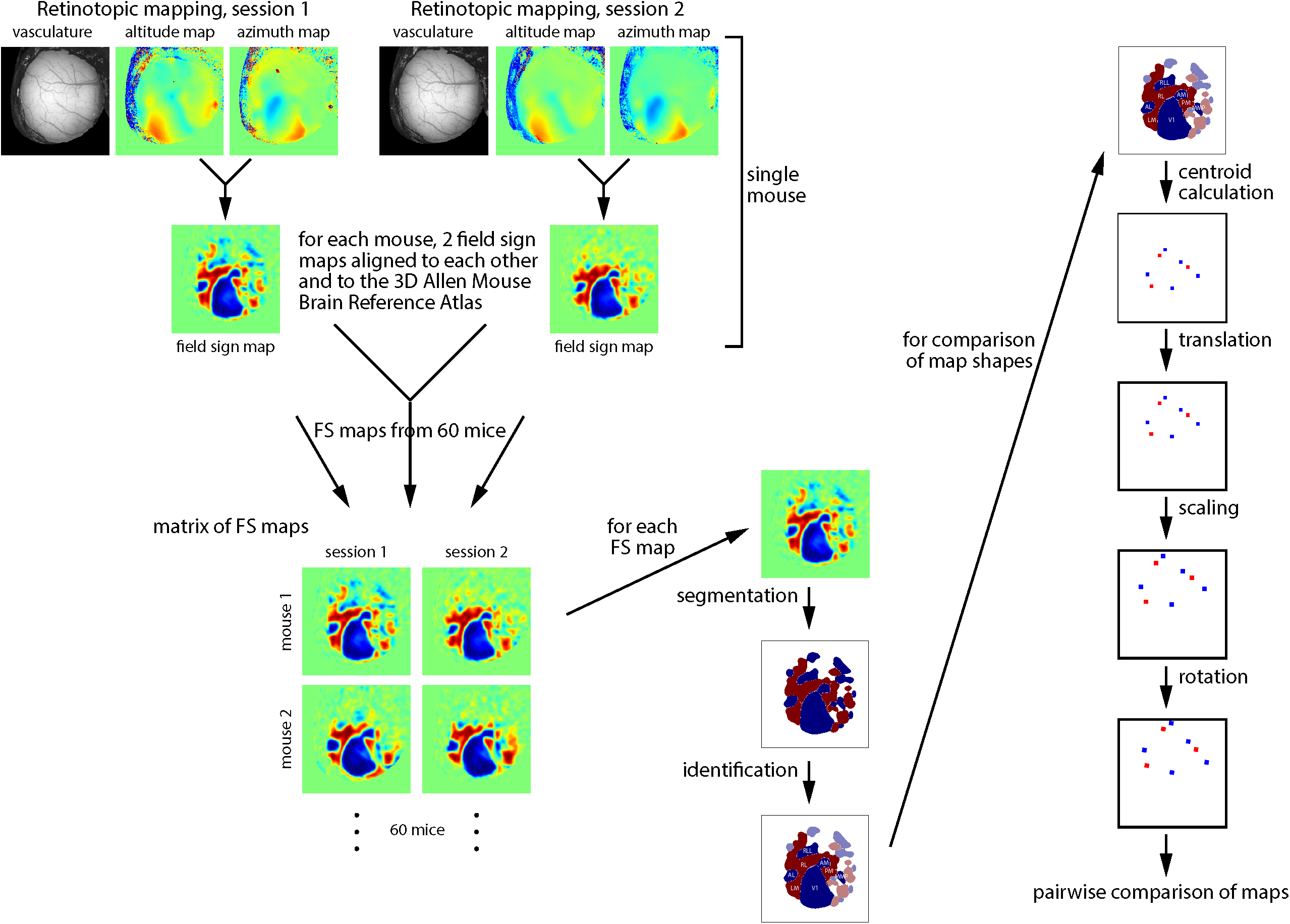
Schematic of analysis workflow. Schematic illustration of the sequence of steps in the core of the analysis.

**Figure 2 - figure supplement 1.**
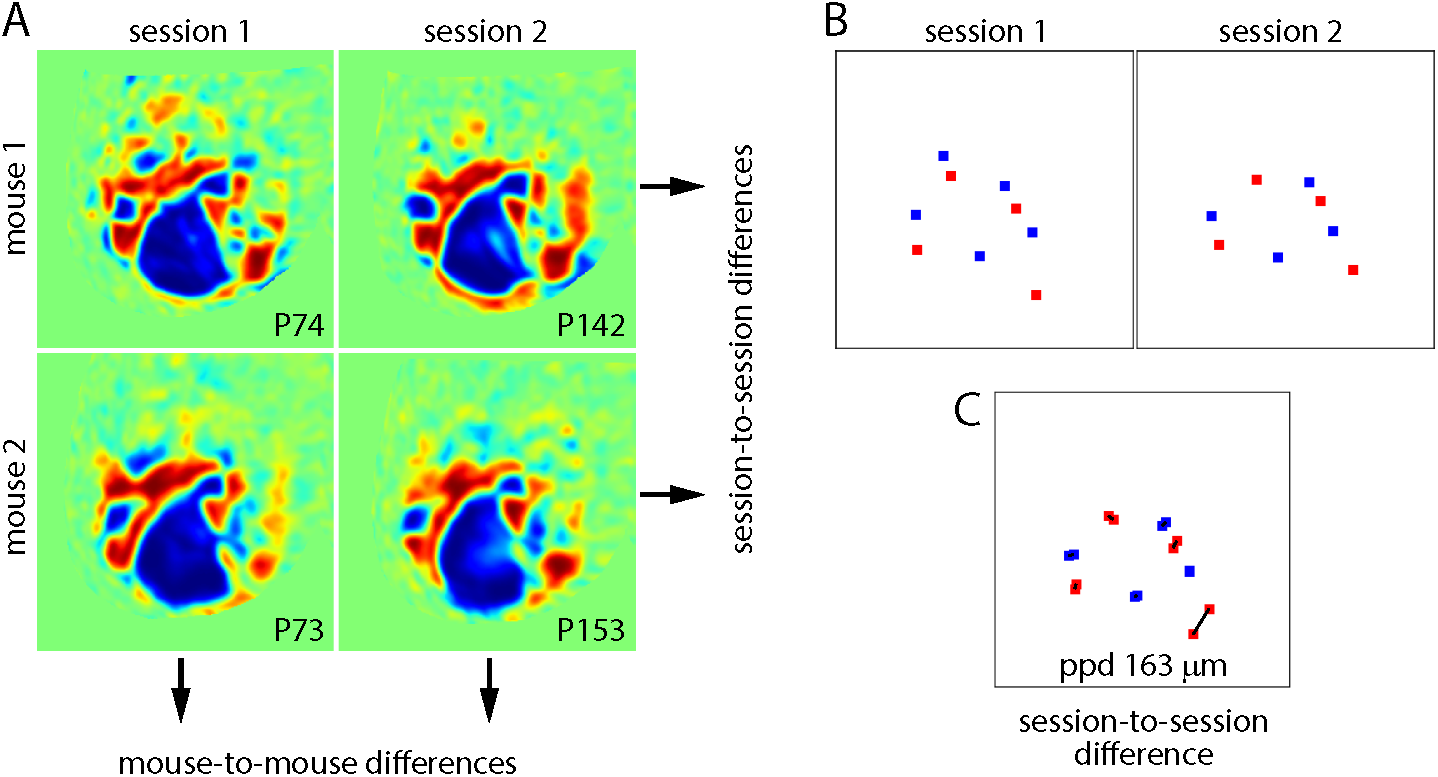
(A) Field sign maps from two mice. For each map, the postnatal age (in days) during imaging is provided. Comparison across mice (across rows) describes the sum of biological variation and measurement noise. Comparison across imaging sessions (down columns) describes measurement noise. The difference between the two comparisons provides an estimate of biological variation. (B) Maps of centroid locations for each field sign patch in mouse 1. Each centroid is colored to match the field sign of its parent field sign patch. (C) Comparison of two maps, with the distances between patches illustrated with black lines. The paired patch distance (ppd) is the mean of these distances.

## MATERIALS AND METHODS

### Mouse lines

Retinotopic maps were generated from 60 mice of 7 genotypes: 20 Cux2-CreERT2;Camk2a-tTA;Ai93 mice, 5 Emx1-IRES-Cre;Camk2a-tTA;Ai93 mice, 1 Emx1-IRES-Cre;Camk2a-tTA;Ai94 mouse, 7 Nr5a1-Cre;Camk2a-tTA;Ai93 mice, 10 Rbp4-Cre;Camk2a-tTA;Ai93 mice, 11 Rorb-IRES2-Cre;Camk2a-tTA;Ai93 mice and 6 Scnn1a-Tg3-Cre;Camk2a-tTA;Ai93 mice.

Mice were crosses of the following lines:

Cux2-CreERT2: Franco *et al.*, 2012; https://www.mmrrc.org/catalog/sds.php?mmrrc_id=32779).

Emx1-IRES-Cre: B6.129S2-Emx1tm1(cre)Krj/J; Gorski *et al.*, 2002; https://www.jax.org/strain/005628)

Nr5a1-Cre: Dhillon *et al.*, 2006; https://www.mmrrc.org/catalog/sds.php?mmrrc_id=34276

Rbp4-Cre_KL100: Gong *et al.*, 2007; https://www.mmrrc.org/catalog/sds.php?mmrrc_id=31125

Rorb-IRES2-Cre: Oh *et al.*, 2014; Harris *et al.*, 2014; https://www.jax.org/strain/023526

Scnn1a-Tg3-Cre: Madisen *et al.*, 2010; https://www.jax.org/strain/009613

CaMK2a-tTA: B6.Cg-Tg(Camk2a-tTA)1Mmay/DboJ; Mayford *et al.*, 1996; https://www.jax.org/strain/007004

Ai93: B6;129S6-Igs7^tm93.1(tetO-GCaMP6f)Hze^/J; Madisen *et al.*, 2015; https://www.jax.org/strain/024103

Ai94: B6.Cg-*Igs7*^*tm94.1(tetO-GCaMP6s)Hze*^/J; Madisen *et al.*, 2015; https://www.jax.org/strain/024104

### Retinotopic mapping and identification of visual areas

Retinotopic maps were generated by intrinsic signal imaging as part of the Allen Brain Observatory data product (http://observatory.brain-map.org/visualcoding/) and imaging methods are described in the Allen Brain Observatory literature (http://help.brain-map.org/display/observatory/Documentation?preview=/10616846/10813483/VisualCoding_Overview.pdf). Retinotopic mapping was performed under isoflurane anesthesia (1-1.4%, inhaled). Altitude and azimuth maps were converted to a field sign map and the borders between areas were identified using code presented in our previous paper (Zhuang *et al.*, 2017). Here we took the additional step of automating the identification of 11 visual areas.

### Alignment of maps

Maps were aligned across mice and imaging sessions. Images from the first imaging session were aligned to the 3D Allen Mouse Brain Reference Atlas as described previously (Kuan *et al.*, 2015; http://help.brain-map.org/display/mouseconnectivity/Documentation?preview=/2818171/10813534/Mouse_Common_Coordinate_Framework.pdf). Images from the second imaging session were aligned to the first session using the surface vasculature images and a custom routine.

### Translation, scaling and rotation of maps

In Procrustean morphometrics, shape is what remains after the removal of differences in translation, scale (or size) and rotation (Goodall, 1991; Webster & Sheets, 2010; Zelditch, Swiderski & Sheets, 2012). To compare the shapes of retinotopic maps (figure 2), we used a Procrustean approach, eliminating map-to-map differences in translation, scale and rotation. We converted each map into a collection of centroids, one for each field sign patch. We translated maps such that the centroid of V1 was at the origin. We normalized the size of each map using the centroids of patches V1, RL and PM since these three patches were identified in every map. These three centroids form a triangle and we scaled each map such that the circumference of this triangle equaled one. Finally, we rotated each map about the origin (the centroid of V1) to minimize the sum of distances between the centroids of corresponding patches (RL in map 1 vs RL in map 2, LM vs LM, etc.).

### Data sets and analysis code

We include two Jupyter notebooks with this manuscript. The ‘data viewer’ notebook provides code to download and view the data sets. The ‘analysis’ notebook contains most of the plots and analyses in the manuscript and some additional analyses.

**Figure 4.**
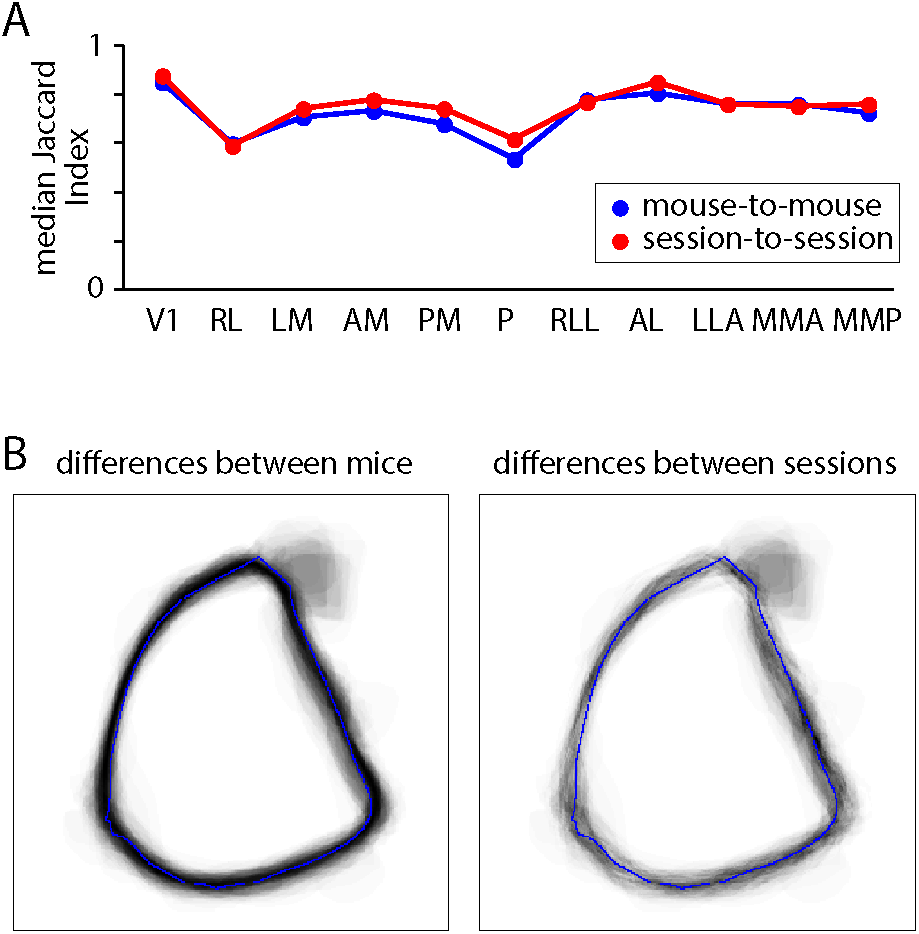
Biological variation in the shapes of field sign patches. (A) Median of the distribution of the Jaccard Indices for mouse-to-mouse and session-to-session comparisons, for each patch. (B) Cumulative differences in V1 for mouse-to-mouse and session-to-session pairwise comparisons.

## ACKNOWLEDGEMENTS

We wish to thank the Allen Institute founder, Paul G. Allen, for his vision, encouragement and support.

